# Use of a glycan library reveals a new model for enteric virus oligosaccharide binding and virion stabilization

**DOI:** 10.1101/834101

**Authors:** Hua Lu, Mark A. Lehrman, Julie K. Pfeiffer

## Abstract

Enteric viruses infect the gastrointestinal tract and bacteria can promote replication and transmission of several enteric viruses. Viruses can be inactivated by exposure to heat or bleach, but poliovirus, coxsackievirus B3, and reovirus can be stabilized by bacteria or bacterial polysaccharides, limiting inactivation and aiding transmission. We previously demonstrated that certain N-acetylglucosamine (GlcNAc)-containing polysaccharides can stabilize poliovirus. However, the detailed virus-glycan binding specificity and glycan chain length requirements, and thus the mechanism of virion stabilization, has been unclear. A previous limitation was our lack of defined-length glycans to probe mechanisms and consequences of virus-glycan interactions. Here, we generated a panel of polysaccharides and oligosaccharides to determine the properties required for binding and stabilization of poliovirus. Poliovirus virions are non-enveloped icosahedral 30 nm particles with 60 copies of each of four capsid proteins, VP1-4. VP1 surrounds the fivefold axis and our past work indicates that this region likely contains the glycan binding site. We found that relatively short GlcNAc oligosaccharides, such as a six unit GlcNAc oligomer, can bind poliovirus but fail to enhance virion stability. Virion stabilization required binding of long GlcNAc polymers of greater than 20 units. Our data suggest a model where GlcNAc polymers greater than 20 units bind and bridge adjacent fivefold axes, thus aiding capsid rigidity and stability. This study provides a deeper understanding of enteric virus-bacterial glycan interactions, which is important for virion environmental stability and transmission.

**Importance:** Enteric viruses are transmitted through the fecal-oral route, but how enteric viruses survive in the environment is unclear. Previously, we found that bacterial polysaccharides enhance poliovirus stability against heat or bleach inactivation, but the specific molecular requirements have been unknown. Here we showed that certain short chain oligosaccharides can bind to poliovirus but do not increase virion stability. Long chain polysaccharides bind and may bridge adjacent sites on the viral surface, thus increasing capsid rigidity and stability. This work defines the unique interactions of poliovirus and glycans, which provides insight into virion environmental stability and transmission.

## Introduction

Enteric viruses primarily infect and replicate in the gastrointestinal tract of the host and are transmitted by the fecal–oral route. Over 100 types of pathogenic enteric viruses are excreted in human and animal feces (Hurst and Gerba 1989). Infections are associated primarily with diarrhea and vomiting in humans and may also cause respiratory infections, hepatitis, and diseases that have high mortality rates, such as encephalitis, and paralysis (Kocwa-Haluch 2001). The picornavirus poliovirus is spread by the fecal-oral route, but can rarely disseminate to the central nervous system, causing paralysis and death (Racaniello 2006).

In the intestine, enteric viruses encounter huge numbers of microbes, collectively known as the microbiota. Intestinal microbiota can promote infection of enteric viruses such as poliovirus, coxsackievirus B3, reovirus, rotavirus, mouse mammary tumor virus, and norovirus through direct effects on virion stability/attachment or indirect effects on host immune responses (Kane, Case et al. 2011; Kuss, Best et al. 2011; Jones, Watanabe et al. 2014; Kernbauer, Ding et al. 2014; Robinson, Jesudhasan et al. 2014; Uchiyama, Chassaing et al. 2014; Baldridge, Nice et al. 2015). For example, using poliovirus as a model enteric virus, we found that poliovirus binds to bacterial surface lipopolysaccharide (LPS) (Kuss, Best et al. 2011; Robinson, Jesudhasan et al. 2014). LPS binding reduced premature genomic RNA release, and increased virion stability and host cell attachment. We found that long N-acetylglucosamine (GlcNAc)-containing polysaccharides can stabilize poliovirus, but a six unit GlcNAc oligosaccharide, GlcNAc6, could not. Recent comparison studies of poliovirus with other members of the *Picornaviridae*, such as coxsackievirus B3, Aichi virus and mengovirus, revealed shared and distinct effects of bacteria on virion stability (Aguilera, Nguyen et al. 2019).

A poliovirus mutant with reduced LPS binding has shed light on potential binding sites on the virion as well as consequences of polysaccharide binding. Poliovirus capsids are icosahedral structures composed of 60 copies of each of four proteins, VP1 to VP4. We previously showed that VP1-T99K mutant poliovirus has reduced binding to LPS and is not stabilized by LPS at physiological temperature (Robinson, Jesudhasan et al. 2014). Deletion of the T99 residue did not influence LPS binding and the T99K mutant phenotype was conditional, with no LPS-mediated stabilization at 37°C, but with wild-type levels of LPS-mediated stabilization at 42°C. These results suggest that VP1-T99 is not directly involved in LPS binding but is likely close to the binding site. Importantly, the LPS low binding T99K mutant virus had a transmission defect in mice when virion instability was a selective pressure (Robinson, Jesudhasan et al. 2014).

Our previous glycan binding studies of poliovirus were performed using bacterial LPS, which is composed of polysaccharide and lipid A (Kuss, Best et al. 2011; Robinson, Jesudhasan et al. 2014). Whether one or both parts of LPS contribute to binding and stabilization of poliovirus is not clear. The polysaccharides of LPS are very diverse in structure, and not all of them contain GlcNAc residues, indicating binding specificity may be not limited to GlcNAc-containing glycans. The relatively large molecular weight and heterogeneity of LPS adds complexity and hinders further characterization of glycan binding avidity and binding sites. Moreover, the glycan chain length requirements for binding/stabilization of poliovirus are unclear.

In this study, we used poliovirus as a model enteric virus and examined the properties of glycans required for binding and stabilization of viral particles. A limitation of prior work was the absence of key reagents to examine effects of glycan chain length on virion binding and stabilization. While our past studies examined virion stability in the presence of dozens of glycans, we lacked glycans of defined lengths in a key range (6-30 unit long oligomers), and our direct binding assays were limited to commercially available biotinylated LPS (Kuss, Best et al. 2011; Robinson, Jesudhasan et al. 2014). Here, we generated a library of different polysaccharides and purified oligosaccharides of different chain lengths and we found that acetylated polysaccharides bind to and stabilize poliovirus. Short chain GlcNAc oligomers could bind but did not stabilize poliovirus. Viral stabilization required GlcNAc oligomers longer than 20 units, likely through bridging of multiple binding sites on virion surface. In conclusion, this work defines the unique interactions of poliovirus and bacterial glycans which may provide insight into virion environmental stability and transmission.

## Results

Previously we found that bacterial LPS can stabilize poliovirus against heat or bleach induced inactivation (Kuss, Best et al. 2011; Robinson, Jesudhasan et al. 2014). LPS contains lipid A and polysaccharide, but whether one or both contributes to poliovirus stabilization is unclear. To investigate this, we used LPS from *E. coli* O127:B8 and performed weak acid hydrolysis (Fig. 1A), which was sufficient to release polysaccharide from lipid A without polysaccharide degradation, as shown by fluorophore-assisted carbohydrate electrophoresis (FACE). FACE is a rapid and sensitive gel electrophoresis method to examine monosaccharide composition and to profile oligosaccharide length. In FACE, a fluorophore is linked to the reducing termini of glycans by reductive amination, followed by gel electrophoresis and detection using ultraviolet light. The identity of each band can be deduced by comparing the migration rate with proper standard. We analyzed the monosaccharide composition of LPS from *E. coli* O127:B8 and confirmed that it mainly contains N-acetyl-galactosamine (GalNAc), galactose, and fucose (Fig.1B), consistent with previous reports (Stenutz, Weintraub et al. 2006). We purified detoxified LPS and lipid A by phase separation and tested their abilities to stabilize poliovirus using an *in vitro* thermal inactivation assay. 10^6^ PFU of poliovirus was incubated with compounds at 45°C for 5 hours, followed by plaque assay on Vero cells to determine the number of remaining infectious particles. We found LPS very efficiently stabilized poliovirus over a wide concentration range (0.01-1 mg/ml) (Fig. 1C), in accordance with previous results (Kuss, Best et al. 2011; Robinson, Jesudhasan et al. 2014). Detoxified LPS (dLPS), which is a polysaccharide usually consisting of 50-100 monosaccharide residues, stabilized poliovirus but was about 10− to 100-fold less effective compared with intact LPS. Lipid A from *E. coli* stabilized poliovirus as well (Fig. 1D). In contrast, 10 different monosaccharides and 4 oligosaccharides did not stabilize poliovirus, even at high concentrations (20 mg/ml). Overall, these data indicate both the polysaccharide and lipid A moieties of LPS contribute to poliovirus stabilization. Polysaccharides, but not monosaccharides or short oligosaccharides, can stabilize poliovirus against heat inactivation.

**Fig 1.**
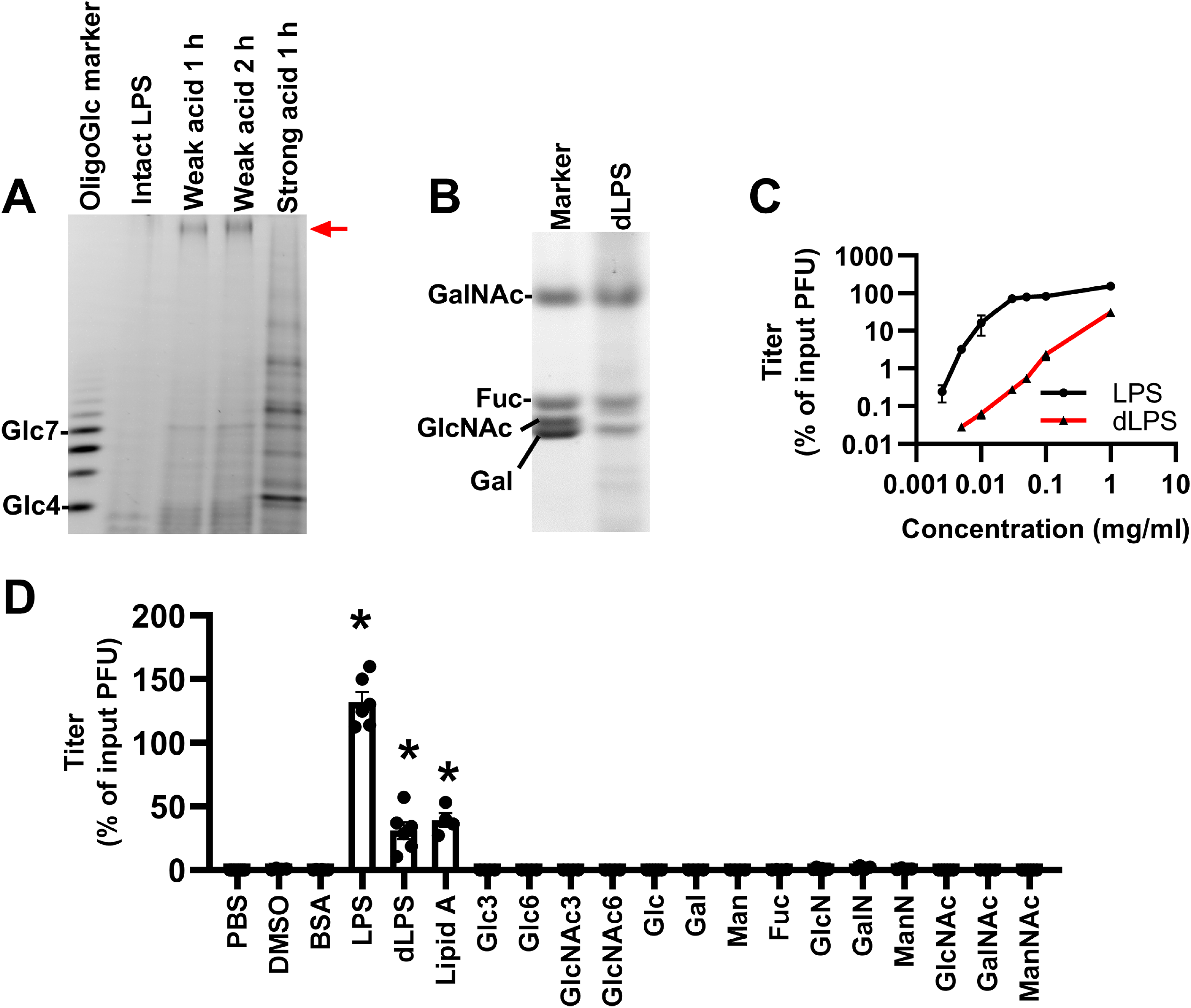
Components of LPS that contribute to poliovirus stabilization. A. LPS from *E. coli* O127:B8 was hydrolyzed by acid and analyzed by FACE. Intact LPS cannot be labeled with fluorophore, and therefore does not generate a band in the FACE gel. Weak acid treatment of LPS (2% acetic acid, 100°C) is sufficient to release intact polysaccharide (detoxified LPS), as indicated by the red arrow. Strong acid treatment of LPS (0.1 M HCl, 100°C) generated multiple degraded fragments of the polysaccharide. Glucose oligomers (Glc4 to Glc7) were used to roughly indicate glycan sizes. B. FACE analysis of the monosaccharide composition of the detoxified LPS from *E. coli* O127:B8 following strong acid hydrolysis. N-acetyl-galactosamine, fucose, and galactose were detected compared with monosaccharide standards. C. Detoxified LPS (dLPS) from *E. coli* O127:B8 stabilizes poliovirus, but at a lower efficiency than intact LPS. 10^6^ PFU of poliovirus was incubated with compounds at 45°C for 5 hours. Remaining titers after incubation were calculated by plaque assays and normalized to untreated control. D. Polysaccharide (dLPS) and lipid components of LPS contribute to poliovirus stabilization, but monosaccharides or short oligosaccharides do not. Concentration of each compound: 1 mg/ml for DMSO, BSA, LPS, dLPS and lipid A; 20 mg/ml for monosaccharides; 5 mg/ml for oligosaccharides. 10^6^ PFU of poliovirus was incubated with compounds at 45°C for 5 hours. Remaining titers after incubation were calculated by plaque assays and normalized to untreated control. Glycan abbreviations: D-glucose (Glc), D-galactose (Gal), D-mannose (Man), L-fucose (Fuc), D-glucosamine (GlcN), D-galactosamine (GalN), D-mannosamine (ManN), N-acetyl-D-glucosamine (GlcNAc), N-acetyl-D-galactosamine (GalNAc), N-acetyl-D-mannosamine (ManNAc), maltotriose (Glc3), maltohexaose (Glc6), tri-N-acetylchitotriose (GlcNAc3), hexa-N-acetylchitohexaose (GlcNAc6). N=4-8, error bars represent mean ± SEM, * p<0.05.

Since the detoxified LPS from *E. coli* O127:B8 contains GalNAc but not GlcNAc, we hypothesized that acetyl groups in polysaccharide are important for binding and stabilization of poliovirus. To test this, we examined the effects of a group of polysaccharides on viral particle thermal stability (Fig. 2A). We found chitin (GlcNAc homopolymer) and peptidoglycan (GlcNAc-containing polysaccharide) stabilized more than 50% of input viruses against heat inactivation. The acetyl groups in chitin can be removed to convert chitin into chitosan. We found that chitosan (85% deacetylated) only stabilized about 5% of input viruses, and more than 99.99% of input viruses were inactivated when coincubated with PBS or cellulose (glucose homopolymer). Randomly introducing O-acetyl groups to cellulose (acetylated cellulose) protected 5% of input viruses against heat inactivation. Thus, acetyl groups in polysaccharide contribute to poliovirus stabilization. We also examined binding of poliovirus with insoluble polysaccharides using ^35^S-labeled virus in a pull-down assay (Fig. 2B). 10^5^ PFU of ^35^S-labeled poliovirus was incubated with insoluble polysaccharides at 37°C for 3 hours. Unbound viruses were washed away with PBS. Polysaccharide-associated viruses were quantified by scintillation counting. Cellulose bound 2.5% of input viruses, comparable to background binding of the inert bead control. The acetylated glycans peptidoglycan and chitin had higher binding capacity (7.5-11%). Similarly, Erickson *et al.* found that *Lactobacillus johnsonii* bacteria with O-acetylated exopolysaccharides (EPS) had more efficient poliovirus binding than isogenic mutant strains that do not produce EPS (Erickson, Jesudhasan et al. 2018). Although chitosan and acetylated cellulose had marginal activity at stabilizing poliovirus, we did not detect any statistically significant binding compared to cellulose in our pulldown assay. Overall, these data suggest acetyl groups in polysaccharides contribute to binding and stabilization of poliovirus but other factors such as carbohydrate conformation and linkage of acetyl groups may also impact binding.

**Fig 2.**
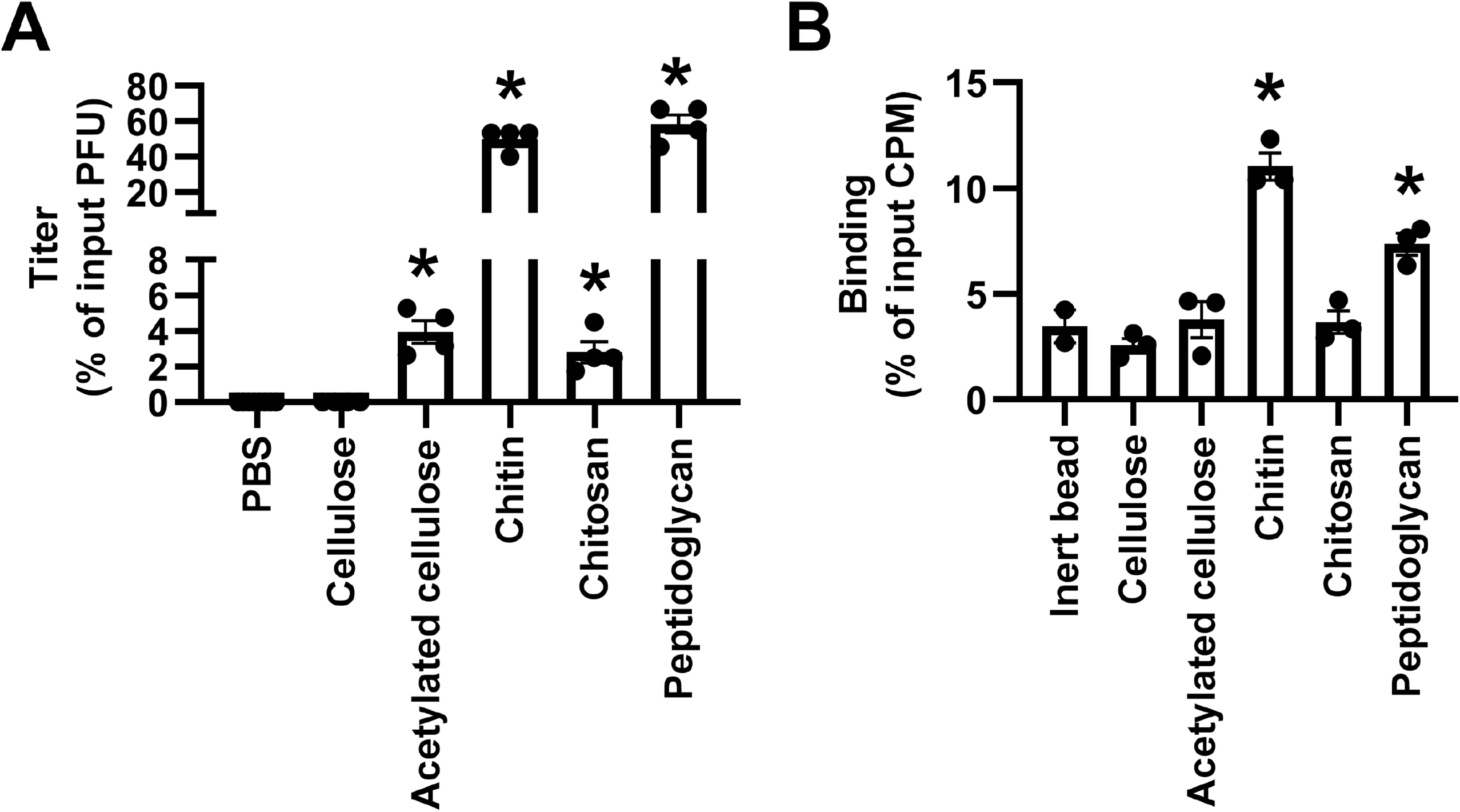
Poliovirus binds to and is stabilized by acetylated polysaccharides. A. Acetylated polysaccharides stabilize poliovirus against *in vitro* thermal inactivation. 10^6^ PFU of poliovirus was incubated with 1 mg/ml of each compound at 45°C for 5 hours. Remaining titers after incubation were quantified by plaque assays and normalized to untreated control. B. Acetylated polysaccharides bind poliovirus in a direct pulldown assay. 10^5^ PFU ^35^S-labeled poliovirus was incubated with 500 μg insoluble polysaccharides. After incubation, samples were centrifuged, washed, and polysaccharide-associated virus in the pellet was measured by scintillation counting. N=2-4, error bars represent mean ± SEM, * p<0.05.

Short oligosaccharides do not stabilize poliovirus, but whether they bind to the virion is unknown. Chitin is a GlcNAc homopolymer and we used it to generate low molecular weight GlcNAc oligosaccharides by acid hydrolysis and we fractionated them by size exclusion chromatography. We obtained GlcNAc oligomers containing 6 to 17 units, as shown by FACE (Fig. 3A). To examine their virion binding abilities, we used lectin cross-linked agarose beads to immobilize oligosaccharides on the bead surface followed by pulldown assay (Fig. 3B). Wheat germ agglutinin (WGA) specifically binds GlcNAc-containing glycans, and was used to bind oligosaccharides to the beads. As a negative control, concanavalin A (Con A) binds glucose-containing glycans. Nonspecific binding sites were blocked with BSA. We used high molar ratios of glycans to ensure that the glycan binding sites on the bead surface were saturated. Excess unbound glycans were washed away with PBS. We coincubated 10^5^ PFU of ^35^S-labeled polioviruses with oligosaccharide-coated beads at 37°C for 3 hours and then measured bead-associated viruses by liquid scintillation counting. We found GlcNAc3-coated WGA beads did not pulldown virus, similar to PBS control. GlcNAc6-coated WGA beads bound 4% of input viruses, while GlcNAc12-14 bound 50% of input viruses, indicating the pulldown efficiency of WGA beads is correlated with the chain length of the coated GlcNAc oligosaccharides. Because WGA binds one to two GlcNAc residues within its glycan binding sites, GlcNAc3-coated WGA beads likely only have 1-2 units exposed on the bead surface, which likely explains the deficiency in the pulldown assay. Although the detoxified LPS of *E. coli* O127:B8 can stabilize poliovirus, it did not pulldown poliovirus here due to lack of GlcNAc residues required for WGA bead binding. The negative control, glucose oligosaccharide coated Con A beads, did not bind poliovirus as well. These data suggest that short GlcNAc oligosaccharides can bind to poliovirus, but they fail to stabilize the virus.

**Fig 3.**
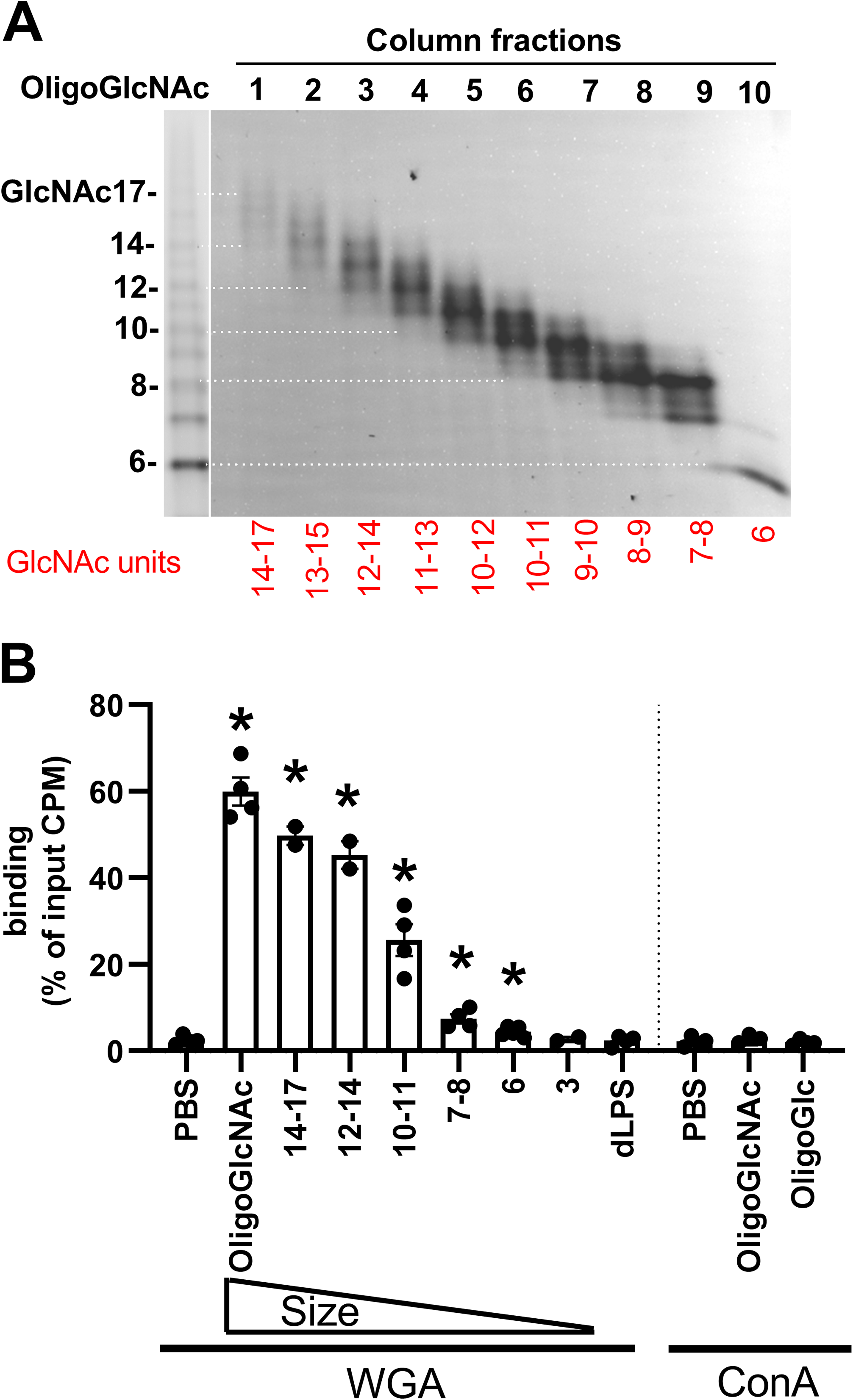
Poliovirus binds low molecular weight GlcNAc oligosaccharides. A. Preparation of low molecular weight GlcNAc oligosaccharides. Chitin was hydrolyzed by acid to generate water soluble oligosaccharides. GlcNAc oligomers were fractionated by Bio-Gel P-4 size exclusion column and analyzed by FACE. The left lane is oligo-GlcNAc standard. B. Poliovirus binds to GlcNAc oligomer coated WGA-agarose beads. 10^5^ PFU of ^35^S-labeled poliovirus was incubated with glycan coated lectin-agarose beads. After incubation, bead-associated virus was measured by scintillation counting. WGA lectin binds GlcNAc-containing glycans; ConA binds glucose-containing glycans. N=2-5, error bars represent Mean ± SEM, * p<0.05.

GlcNAc6 binds but does not stabilize poliovirus, suggesting it may act as a competitive inhibitor of polysaccharide-mediated virion stabilization. To test this, we coincubated poliovirus with polysaccharides in the presence or absence of excess GlcNAc6 in our *in vitro* thermal inactivation assay (Fig. 4A). We found coincubation of 1 mg/ml GlcNAc6 with 0.05 mg/ml detoxified LPS reduced the recovered titer compared with detoxified LPS alone. Similarly, 1 mg/ml GlcNAc6 inhibited 0.5 mg/ml chitin-mediated stabilization of poliovirus, suggesting that GlcNAc6 competes with long polysaccharides for the limited glycan binding sites on virion surface. However, we did not observe any titer decrease when poliovirus was coincubated with 0.01 mg/ml LPS in the presence of 1 mg/ml GlcNAc6. The lack of competition of GlcNAc6 for intact LPS may be due to the virion stabilizing effects of lipid A. Control Glc6 had no effect on polysaccharide-mediated stabilization of poliovirus. The effects of GlcNAc6 were concentration dependent, with inhibition requiring at least 10-fold excess GlcNAc6 by weight (Fig. 4B). Since excess GlcNAc6 decreased dLPS-mediated virion stabilization, we wondered whether shorter GlcNAc oligomers would have activity. We performed acid hydrolysis of the long GlcNAc polysaccharide chitin, purified short GlcNAc oligomers using size exclusion chromatography, and analyzed them by FACE (Fig. 4C). Poliovirus was exposed to 0.05 mg/ml dLPS in the presence or absence of 1 mg/ml GlcNAc oligomers 3, 4, 5, 6, or 7 units long in our *in vitro* thermal inactivation assay (Fig. 4D). We found that inhibition of dLPS-mediated stabilization became stronger as the glycan chain length increased from 3 to 6 GlcNAc residues. GlcNAc7 did not cause further titer reduction. Overall, these data indicate that excess short GlcNAc oligosaccharides inhibit polysaccharide-mediated stabilization of poliovirus, suggesting that GlcNAc oligomers as short as 5 units may bind poliovirus but fail to stabilize.

**Fig 4.**
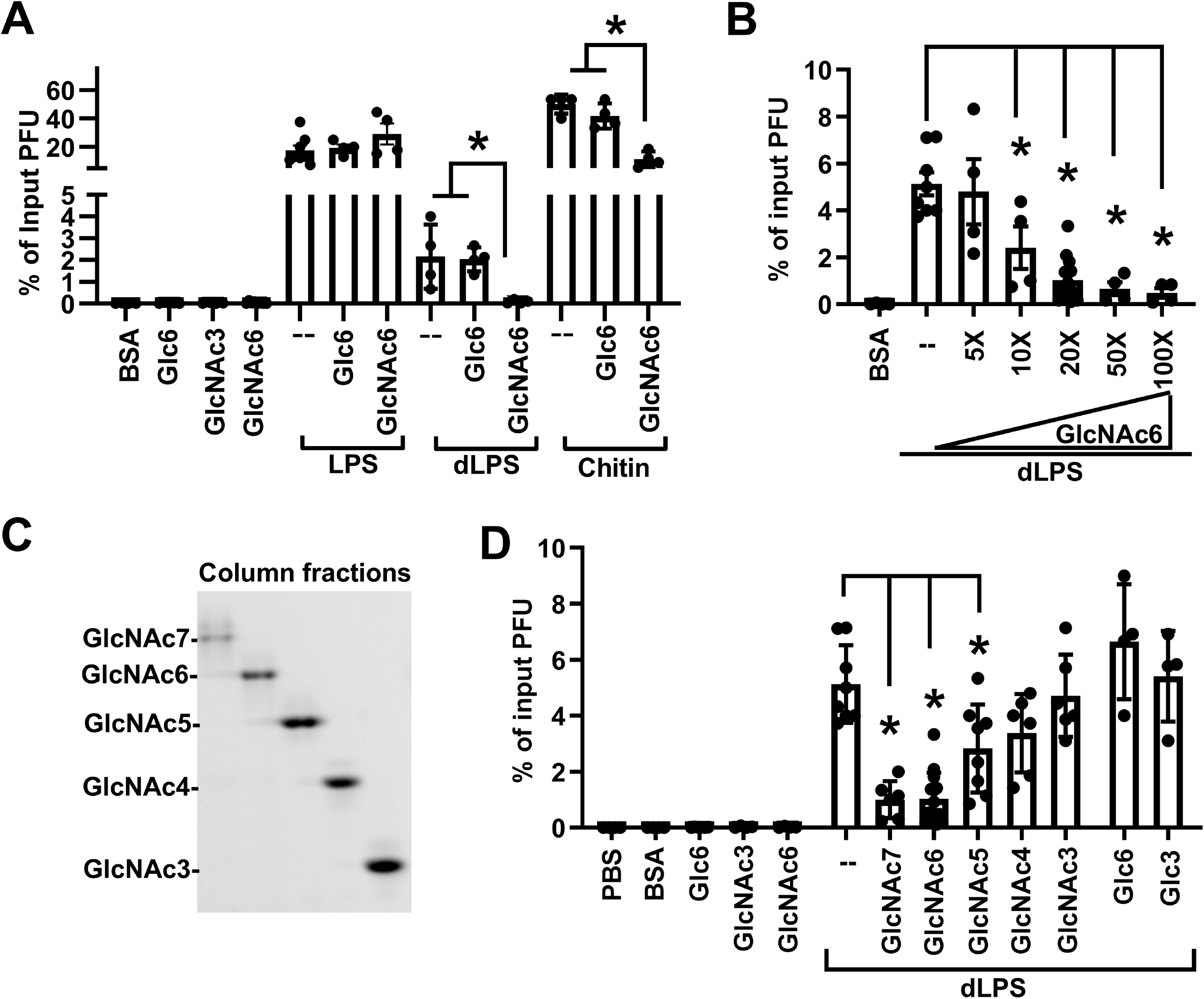
Excess GlcNAc6 inhibits polysaccharide-mediated stabilization of poliovirus. A. Excess GlcNAc6 inhibits detoxified LPS (dLPS) and chitin-mediated stabilization of poliovirus. 10^6^ PFU of poliovirus was incubated with compounds at 45°C for 5 hours. Remaining titers after incubation were quantified by plaque assay and normalized to untreated control. N=4-8. B. Inhibition effect of GlcNAc6 is concentration dependent. Experiments were performed as in (A), but different concentrations of GlcNAc6 were used. N=4-14. C. FACE analysis of purified glycans containing 3 to 7 GlcNAc units. Chitin was hydrolyzed by acid to generate water soluble oligosaccharides. GlcNAc oligomers were fractionated by Bio-Gel P-4 size exclusion column and analyzed by FACE. D. The chain lengths of GlcNAc oligomers affect their inhibition efficiency. Experiments were performed as in (A). Concentration of each compound: 0.01 mg/ml for LPS; 0.05 mg/ml for dLPS; 0.5 mg/ml for chitin; 1 mg/ml for BSA, GlcNAc3, GlcNAc4, GlcNAc5, GlcNAc6, GlcNAc7, Glc3 and Glc6. N=4-14. Error bars represent Mean ± SEM, * p<0.05.

To define the minimum glycan length requirement for stabilization, we examined activity of GlcNAc oligomers of a variety of lengths in titer-based virion stability assays and in a virion RNA release assay. We performed acid hydrolysis of chitin, purified high molecular weight GlcNAc oligomers using size exclusion chromatography, and analyzed them by FACE. Due to limitations of FACE gel resolution, glycans longer than 20 GlcNAc residues appear as a smear in the top of the gel whereas glycans containing less than 20 GlcNAc residues are well separated (Fig. 5A). We performed *in vitro* thermal inactivation experiments by incubating 10^6^ PFU of poliovirus with 0.5 mg/ml of each column fraction at 45°C for 5 hours followed by viral titer assay. We found that fractions 1 and 2, which contained long chain oligosaccharides (>20 GlcNAc residues), were the most effective at stabilizing poliovirus, comparable to intact chitin (Fig. 5B). Fractions 3 and 4 were mixture of short and long glycans (14 to >20 residues), and they also stabilized poliovirus at lower efficiency. Fractions 5 to 10, containing glycans less than 20 GlcNAc residues, did not stabilize poliovirus. We directly measured poliovirus stability using a technique independent from the plaque assay: particle stability thermal release assay (PaSTRY) (Walter, Ren et al. 2012) (Fig. 5C–D). PaSTRY quantifies virion stability by monitoring the temperature at which virion RNA is released. Poliovirus was mixed with SYBR Green II dye and heated in a real-time instrument on a stepwise temperature gradient with fluorescence monitoring. We found that viral RNA release temperature in the presence of PBS was about 45°C, in line with previous results (Walter, Ren et al. 2012). Pre-exposure of poliovirus with fraction 1 or fraction 2, which contain GlcNAc oligomers >20 units, at 37°C for 1 hour increased the RNA release temperature by 2 degrees. Coincubation with Fraction 5, which contains GlcNAc oligomers ~13-18 units, before PaSTRY did not change RNA release temperature, and was similar to the PBS control. Collectively, these results indicated that glycans containing more than 20 GlcNAc residues can stabilize poliovirus.

**Fig 5.**
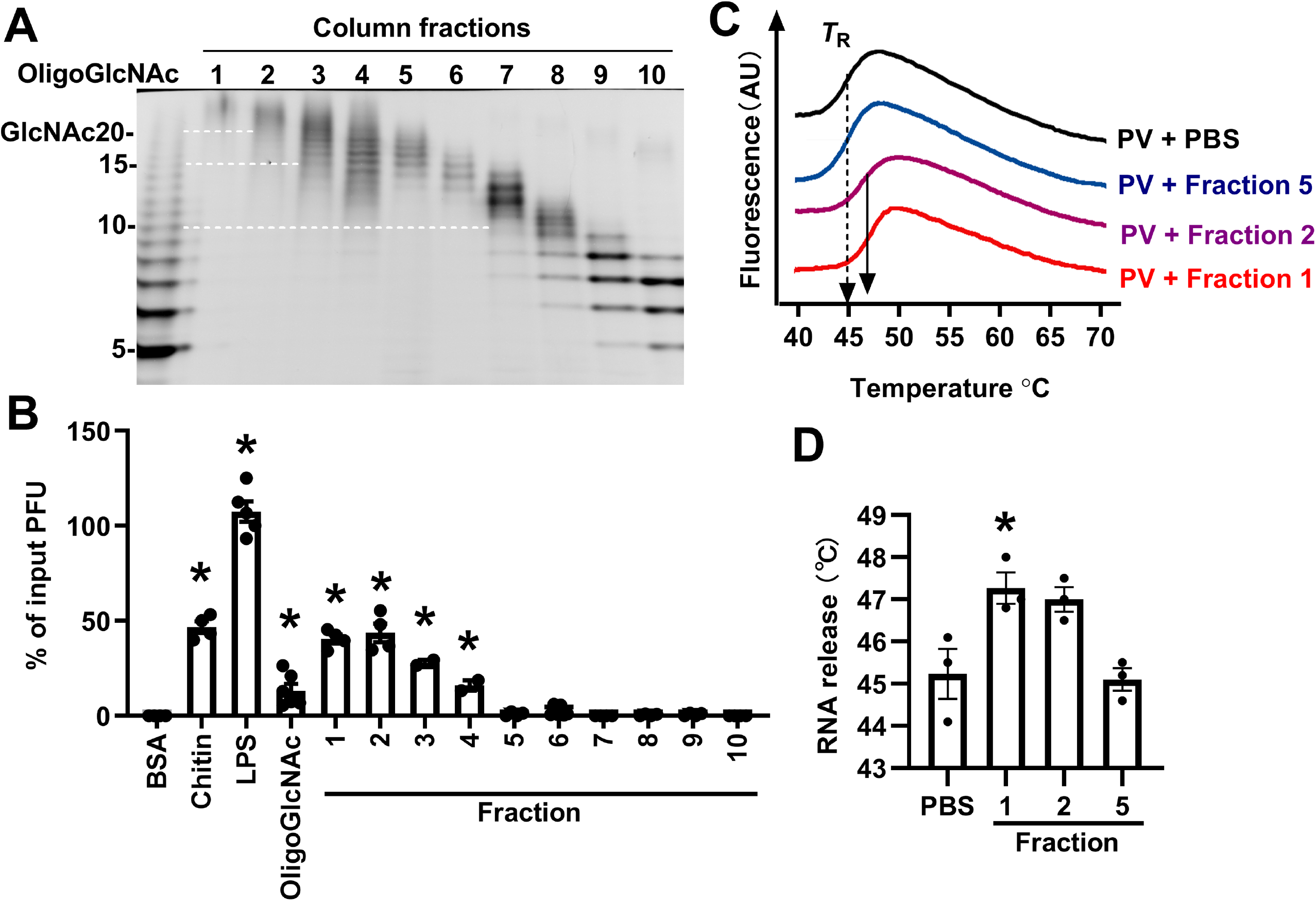
High molecular weight GlcNAc oligosaccharides (>20 units) stabilize poliovirus. A. FACE analysis of Bio-Gel P-10 column fractionated high molecular weight GlcNAc oligosaccharides. B. GlcNAc oligomers longer than 20 units stabilize poliovirus against heat inactivation. 10^6^ PFU of poliovirus was incubated with 0.5 mg/ml of each compound at 45°C for 5 hours. Remaining titer after incubation was quantified by plaque assay and normalized to untreated control. N=2-6. C. Representative profiles of poliovirus particle stability thermo release assay (PaSTRy). Poliovirus was preincubated with 0.5 mg/ml compound for 1 hour at 37°C. Subsequently, viruses were mixed with SYBR green and heated in a real-time machine over a stepwise temperature gradient. The intensity of SYBR green fluorescence was plotted over increasing temperature. Virion RNA release temperature (*T*_R_) represents the temperature causing the highest instantaneous rate of fluorescence change, and is indicated by arrow. AU, arbitrary units. D. Quantification of RNA release temperatures measured by PaSTRy experiments. N=3. Error bars represent Mean ± SEM, * p<0.05.

## Discussion

Enteric viruses are in close contact with bacteria in the gastrointestinal tract and previous studies found that bacteria promote enteric virus replication and transmission (Pfeiffer and Virgin 2016; Robinson 2019; Roth, Grau et al. 2019). Using poliovirus as a model system, we previously demonstrated that binding of bacteria or bacterial polysaccharides can prevent premature release of viral RNA, which is important for the stability of virus in the environment and may aid transmission to the next host. For example, a viral mutant with reduced LPS binding, VP1-T99K, had a transmission defect in mice due to virion instability in feces (Robinson, Jesudhasan et al. 2014). Despite the importance of bacteria-virus interactions, the binding specificity and how glycans bind and stabilize enteric viruses are unclear. The major goal of this study was to determine the glycan requirements for binding and stabilization of poliovirus.

We found that both polysaccharide and lipid components of LPS contribute to poliovirus stabilization (Fig. 1). Poliovirus capsid protein VP1 has a hydrophobic pocket which is occupied by a fatty acid-like molecule, the so-called “pocket factor” (Filman, Syed et al. 1989). Antiviral agents can replace the pocket factor and bind to the hydrophobic pocket, which can stabilize the virus and prevent uncoating (Fox, Otto et al. 1986). A previous study reported increased poliovirus stability in the presence of fatty acids (Dorval, Chow et al. 1989). Unlike the diverse structures of bacterial LPS glycan moieties, the lipid A structure in general is highly conserved, and contains 6 acyl chains linked to the disaccharide backbone (Steimle, Autenrieth et al. 2016). It is possible that one of the acyl chains of lipid A can insert into the hydrophobic pocket and stabilize poliovirus.

The glycan binding specificity of poliovirus is not restricted to GlcNAc containing polysaccharides, but rather acetylated glycans. Acetylated glycans are frequently found in bacterial surface polysaccharides and mammalian intestinal mucin glycans. Glycans can be acetylated through an amide bond (N-acetylation) or ester bond (O-acetylation). Glycan acetylation can alter biological properties of bacteria-host interactions, such as bacteria virulence, disease pathogenesis as well as the host innate and adaptive immune responses (Berti, De Ricco et al. 2018). Moreover, acetylation is a reversible modification. Some bacteria encode deacetylating enzymes which can tune the acetylation level of surface polysaccharides according to environment stress (Benachour, Ladjouzi et al. 2012). Interactions of poliovirus with both N-acetylated (such as chitin and peptidoglycan) and O-acetylated polysaccharides (such as *Lactobacillus johnsonii* EPS) suggest that modulation of the acetylation level of bacterial polysaccharides may impact poliovirus.

We examined binding of poliovirus with insoluble polysaccharides via direct pull-down assays (Fig. 2) and purified GlcNAc oligosaccharides via lectin bead pull-down assays (Fig. 3). We found that long, insoluble GlcNAc-containing polysaccharides such as chitin and peptidoglycan bound poliovirus. Furthermore, short GlcNAc oligomers immobilized on lectin agarose beads bound poliovirus whereas glucose-containing glycans did not. For the GlcNAc oligomers, viral binding efficiency was proportional to glycan length, with 7-8 unit long oligomers binding weakly, but 14-17 unit long oligomers binding efficiently (Fig. 3B). Interestingly, GlcNAc6 had minimal viral binding in the lectin bead pull down assay (Fig. 3B), but was able to competitively inhibit dLPS-mediated viral stabilization (Fig. 4A, B, D). These results suggest that our direct binding assays may be stringent and underestimate binding of short glycans. Regardless, cumulatively, our data show that relatively short GlcNAc oligomers can bind to poliovirus.

Although short GlcNAc glycans could bind to poliovirus, we found that virion stabilization required long GlcNAc polymers (>20 units): Why do short glycans bind virions, but not stabilize them? Poliovirus capsids contain 60 copies of each capsid protein, VP1 to VP4. VP1 surrounds the fivefold axis, and defective LPS binding of the VP1-T99K mutant suggests that this region contains the glycan binding site (Robinson, Jesudhasan et al. 2014). We measured the distance of T99 residues on virion surface using Chimera software based on a poliovirus structure at 3.0 Å resolution (Miller, Hogle et al. 2001; Pettersen, Goddard et al. 2004). We found the distance between closest T99 residues within a fivefold axis is 2.9 nm, which approximately matches the length of GlcNAc6 (3 nm) (Nakamura, Okazaki et al. 2018)(Fig. 6). Conversely, the distance between a T99 residue at one fivefold axis and a T99 residue at an adjacent fivefold axis is 13.4 nm, which approximately matches the length of GlcNAc27 (13.5 nm) (Fig. 6). Thus, we propose a model where short oligomers such as GlcNAc6 are long enough to bind two adjacent sites within a fivefold axis, but this is insufficient to stabilize the virus; conversely, long oligomers (>13.4 nm) bind to and bridge two adjacent fivefold axes, thus aiding structural rigidity and stability of the capsid. This model would explain why excess GlcNAc6 inhibits stabilization by long glycans. We propose that long glycans form chain-like structures on the virion surface and restrain heat induced virion conformation changes that precede premature RNA release.

**Fig 6.**
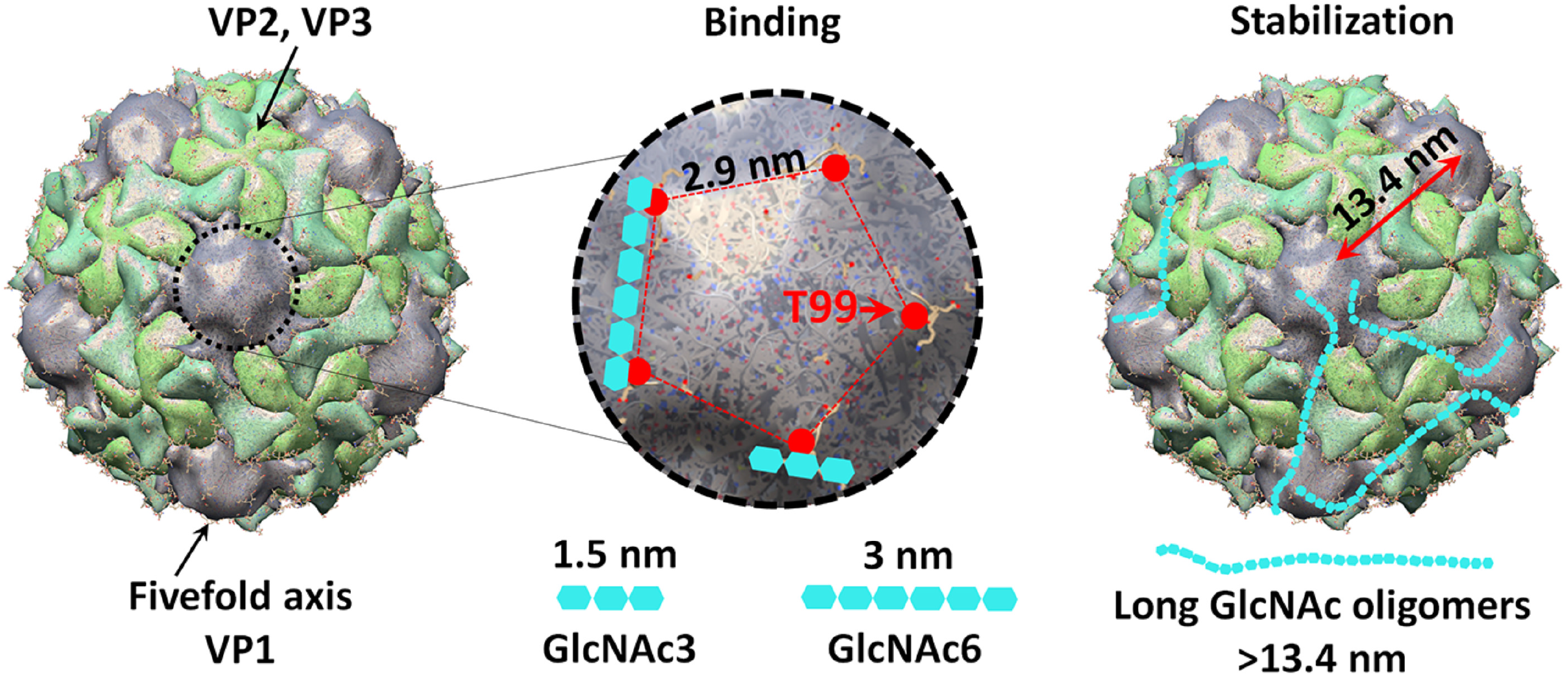
Model of glycan binding and stabilization of poliovirus. The poliovirus capsid contains 60 copies each of VP1, VP2, VP3, and VP4. VP4 is internal and not surface exposed. VP1 surrounds the five-fold symmetry axis, whereas VP2 and VP3 alternate around the three-fold axis. Based on the number of capsid proteins present, each virion may have 60 glycan binding sites. The exposed T99 residue of VP1 is important for polysaccharide binding, and is likely near the glycan binding site. The distance between two nearby T99 residues is 2.9 nm, corresponding with the length of GlcNAc6 (3 nm). Our data suggest that GlcNAc6 can bind two adjacent glycan binding sites within a five-fold axis, resulting in increased avidity compared with monovalent non-binding GlcNAc3 (1.5 nm). Long chain polysaccharides (>13.4 nm) are long enough to potentially bridge nearby fivefold axes and form cage-like structures on the virion surface, and stabilize poliovirus against heat induced conformational change causing premature RNA release.

The glycan binding specificity of poliovirus may not be conserved among other enteric viruses. Human rotavirus and norovirus interact with polymorphic human histo-blood group antigens (HBGAs), and this interaction is thought to be important for infection (Tan and Jiang 2014). Coxsackievirus B3 is more sensitive to microbiota perturbation than poliovirus (Robinson, Acevedo et al. 2019), indicating it may have distinct glycan binding specificity. Bacterial LPS and peptidoglycan also increase reovirus thermostability (Berger, Yi et al. 2017), but the detailed glycan binding specificity is unclear.

Proviral effects of microbiota have been observed for viruses in four different families (Pfeiffer and Virgin 2016). Antibiotic treatment causes microbiota imbalances and can cause antibiotic resistance (Becattini, Taur et al. 2016), making perturbation of the microbiota unviable for treatment of enteric virus infection. As an alternative, targeting the binding of enteric virus to bacterial glycans may provide a promising approach for control of enteric virus infection. Short GlcNAc oligosaccharides can inhibit polysaccharide-mediated poliovirus stabilization. The concentration of bacterial polysaccharide present in the gastrointestinal tract is estimated in the milligram/milliliter range (Bates, Akerlund et al. 2007). Due to such a high concentration, oral administration of short GlcNAc oligosaccharides is not likely to have any anti-viral effects. However, it is useful to characterize glycan-virion binding, which may aid the design of high binding small molecules as novel antiviral drugs. Overall, our work has provided new insight into glycan-virus interactions that impact viral stability.

## Material and Methods

### Virus

Poliovirus work was performed in BSL2+ areas in accordance with practices recommended by the World Health Organization. Poliovirus (serotype 1 virulent Mahoney) cell culture infections and plaque assays were performed using HeLa and Vero cells, respectively (Kuss, Best et al. 2011).

### Acid hydrolysis of LPS

LPS from *E. coli* O127:B8 (L3129, Sigma-Aldrich) was dissolved in 2% acetic acid at 5 mg/ml. After incubation at 100°C for 2 hours with occasional shaking, chloroform and methanol were added to yield final volume ratio of C:M:W=2:1:3. The mixture was vortexed and centrifuged at 5,000 *g* for 10 mins. The resulting upper phase containing detoxified LPS and the lower phase containing Lipid A were collected separately and dried in a Speed-Vac. For monosaccharide analysis, detoxified LPS was dissolved in 1 M HCl and heated at 100°C for 30 mins. Samples were dried in a Speed-Vac.

### FACE analysis

Detoxified LPS, LPS monosaccharides and chitin oligosaccharides were analyzed by FACE (Gao and Lehrman 2006; Gao, Holmes et al. 2013). Briefly, monosaccharides were labeled with 2-aminoacridone (06627, Sigma-Aldrich) and oligosaccharides were labeled with 7-amino-1,3-naphthalenedisulfonic acid (81529, AnaSpec) prior to electrophoresis on 20% acrylamide gels. Gel images were acquired with a UVP Chemidoc-It II Scanner (Analytik Jena, Jena, Germany).

### Preparation and purification of GlcNAc oligomers

Chitin oligomers were prepared via chitin hydrolysis in concentrated hydrochloric acid followed by acetone precipitation (Kazami, Sakaguchi et al. 2015). 2 g chitin (C9752, Sigma) was dissolved in 80 ml concentrated HCl and incubated at 40°C for 1 hour. Samples were added into 1100 ml acetone and stirred overnight at 4°C. The glycan pellet was collected by centrifugation at 5,000 *g* for 10 mins. The pellet was washed twice with acetone, dried at 43°C in an oven, and then 200 ml water was added and stirred overnight at 4°C. Insoluble polysaccharides were removed by centrifugation at 12,000 *g* for 10 mins. The supernatant containing soluble chitin oligosaccharides was loaded on a Dowex AG® 50W-X8 hydrogen form cation exchange column to eliminate any deacetylated products. The flow through fraction was concentrated in a Speed-Vac. The concentrations of oligosaccharides were measured by absorbance at 210 nm using GlcNAc6 as a standard.

For purification of low molecular weight GlcNAc oligomers, a hand-packed Bio-Gel P-4 size exclusion column (1.0 × 50 cm, 1504128, Bio-Rad) was used for glycan separation. Samples were eluted in water at 0.1 ml/min and fractionated 5 mins/tube. For purification of high molecular weight GlcNAc oligomers, a hand-packed Bio-Gel P-10 size exclusion column (1.5 × 50 cm, 1504144, Bio-Rad) was used. Samples were eluted in 20 mM acetic acid at 0.33 ml/min and fractionated 10 mins/tube. Glycans in each fraction were quantified by measuring absorbance at 210 nm and analyzed by FACE.

### ^35^S-labeled poliovirus pulldown assay

^35^S-labeled polioviruses were generated by growing virus in the presence of ^35^S-labeled methionine/cysteine and purification by CsCl gradient ultracentrifugation (Kuss, Best et al. 2011). For insoluble polysaccharide pulldown assay, 5,000 CPM of ^35^S-labeled polioviruses (10^5^ PFU) was mixed with 500 μg polysaccharides in 0.5 ml PBS and incubated at 37°C for 3 h. Polysaccharides were pelleted at 1,500 *g* for 2 mins and washed with 1 ml PBS for three times. Polysaccharide-associated virus was measured by scintillation counting. Peptidoglycan (77140, Sigma) was further purified by proteinase K digestion and PBS wash. All other insoluble polysaccharides were from Sigma without further purification. Inert beads (Dynabeads, 65305, Thermo Fisher) were used as a background binding control.

For binding of low molecular weight GlcNAc oligomers, agarose bead cross-linked lectin (AL-1023, Vector Laboraties) was used to immobilize oligosaccharides on the bead surface. 10 μl bead slurry was incubated overnight with 20 μg glycans in PBS containing 4% BSA. Beads were washed three times to remove unbound glycans and mixed with ^35^S-labeled poliovirus following the same protocol as insoluble polysaccharides. Con A-agarose beads (AL-1003, Vector Laboratories) were used as negative control.

### Poliovirus *in vitro* thermal stability assay

CsCl-gradient purified polioviruses were used for thermal stability experiments. For *in vitro* thermal inactivation, 10^6^ PFU of poliovirus was mixed with compounds at indicated concentrations and incubated at 45°C for 5 h. After incubation, plaque assays were performed in Vero cells and remaining PFU was compared with input to calculate % of input PFU. D-glucose (Glc), D-galactose (Gal), D-mannose (Man), L-fucose (Fuc), D-glucosamine (GlcN), D-galactosamine (GalN), D-mannosamine (ManN), N-acetyl-D-glucosamine (GlcNAc), N-acetyl-D-galactosamine (GalNAc), N-acetyl-D-mannosamine (ManNAc), maltotriose (Glc3), and maltohexaose (Glc6) were purchased from Sigma. Tri-N-acetylchitotriose (GlcNAc3) and hexa-N-acetylchitohexaose (GlcNAc6) were from Caymanchem. The identity of these glycans were confirmed by FACE.

For PaSTRY, 1 μg of poliovirus was preincubated with compounds for 1 h at 37°C to promote binding. After incubation, 10X final concentration of SYBR Green II and buffer (10 mM HEPES, 200 mM NaCl, pH8.0) were added to a final volume of 30 μl (Walter, Ren et al. 2012). Samples were heated in an ABI 7500 real-time instrument from 25°C to 99°C on a 1% stepwise gradient with fluorescence monitoring.

### Statistics

Two-tailed Student’s t tests were used when two groups were compared. p < 0.05 was considered statistically significant.

## Acknowledgements

We thank Broc McCune for helpful comments on the manuscript. This work was supported by a Burroughs Wellcome Fund Investigator in the Pathogenesis of Infectious Diseases award, Howard Hughes Medical Institute Faculty Scholar Award, and R01 AI74668 to JKP.

